# Balancing sensitivity and specificity in preclinical research

**DOI:** 10.1101/2022.01.17.476585

**Authors:** Meggie Danziger, Anja Collazo, Ulrich Dirnagl, Ulf Toelch

## Abstract

The success of scientific discovery in preclinical research is based on the different roles of exploration and confirmation. Exploration involves identifying potential effects (high sensitivity), which are then tested more rigorously during confirmation (high specificity). Here, we examine different experimental strategies and their ability to balance sensitivity and specificity to identify relevant effects. In simulations based on empirical data, we specifically compare a conventional *p*-value based approach with a method based on an *a priori* determined smallest effect size of interest (SESOI). Using a SESOI increases transition rates from exploration to confirmation and leads to higher detection rates across the trajectory. In particular, specificity in the SESOI trajectory increases if number of true effects are low. We conclude that employing a SESOI is superior to a *p*-value based approach in many contexts. Based on our findings, we propose a reconsideration of planning and conducting preclinical experiments, especially when the prior probability of true hypotheses is low.

## Main

Preclinical research is essential for identifying promising therapeutic interventions and to generate robust evidence to support their translation to humans. To achieve these goals, experiments are conducted in two different operating modes.^1^ Early-stage preclinical experiments are *exploratory* with the aim to discover potentially effective interventions and generate hypotheses. These are tested at a later stage under more strict conditions in *confirmatory* mode.^2^

This sequential approach putatively increases the likelihood that generated evidence is robust and potentially reduces translational failures.^3–5^ But which exploratory results are promising and worthwhile to confirm? Here we compare two approaches to select and conduct confirmatory studies with regard to their success in discovering true effects. True here refers to effects with a biologically and/or clinically meaningful magnitude. In one approach, decisions for selecting a study are based on the *p*-value, the ubiquitous criterion in the biomedical publication record. An alternative approach focuses on an *a priori* defined smallest effect size of interest (SESOI), similar to a minimally clinically important difference in clinical trials.^6^

The effectiveness of any approach is based on the different roles exploration and confirmation fulfill in scientific discovery. Via sensitive tests exploration detects potentially true effects among many hypotheses that often have a low prior probability of being true. The sensitivity of a test gives the probability of correctly identifying true effects (also known as the true positive rate or power). As more sensitive criteria invite more false positive results, confirmation must aim at reducing false positives to ensure that only true effects are carried forward to subsequent testing. Specificity, on the other hand, is the ability of a test to correctly reject null effects. The complement of sensitivity is the false negative rate, i.e., the more sensitive a test is the fewer true effects are missed. Likewise, the false positive rate is complementary to specificity. In the null hypothesis significance testing (NHST) framework, for example, the specificity is 95%, if we allow a false positive rate of 5% under the assumption that the null hypothesis is true. In any screening process, sensitivity and specificity have to be weighed against each other^7^ as the two are inversely linked, i.e., raising sensitivity entails a drop in specificity and vice versa.^8^ To complicate matters, ethical, time, and budget constraints limit degrees of freedom in experimental design. Consequently, to prevent false negative results in exploration and to reduce false positives during confirmation, it is necessary to devise strategies to optimize these complementary goals when advancing from exploration to confirmation. Specifically, the likelihood of false negative and false positive outcomes needs to be balanced with the goal to minimize the number of animals needed.

To this end, we simulated two preclinical research trajectories each comprising an exploratory study and a subsequent confirmatory study (Fig. 1a). We based our simulations on a published effect size distribution derived from empirical research^9^ (Fig. 1b; effect sizes are expressed as the standardized difference in means, Hedges’ *g* or Cohen’s *d*). After an initial exploratory study, a decision criterion selected experiments to advance from exploratory to confirmatory mode. One trajectory (Standard) employed the conventional significance threshold (*α* = .05) for this decision using a two-sided two-sample Welch’s *t*-test. The second trajectory (SESOI) used a more lenient threshold based on an *a priori* defined smallest effect size of interest. For this, we estimated the effect size of each exploratory study and its 95% confidence interval (CI). We examined whether this CI covered our SESOI (Hedges’ *g* of 0.5 and 1.0, respectively) and whether the effects were in the hypothesized direction (>0). Importantly, we did not consider significance, thus, with this method significant and non-significant results were carried forward to confirmation.

**Figure 1:**
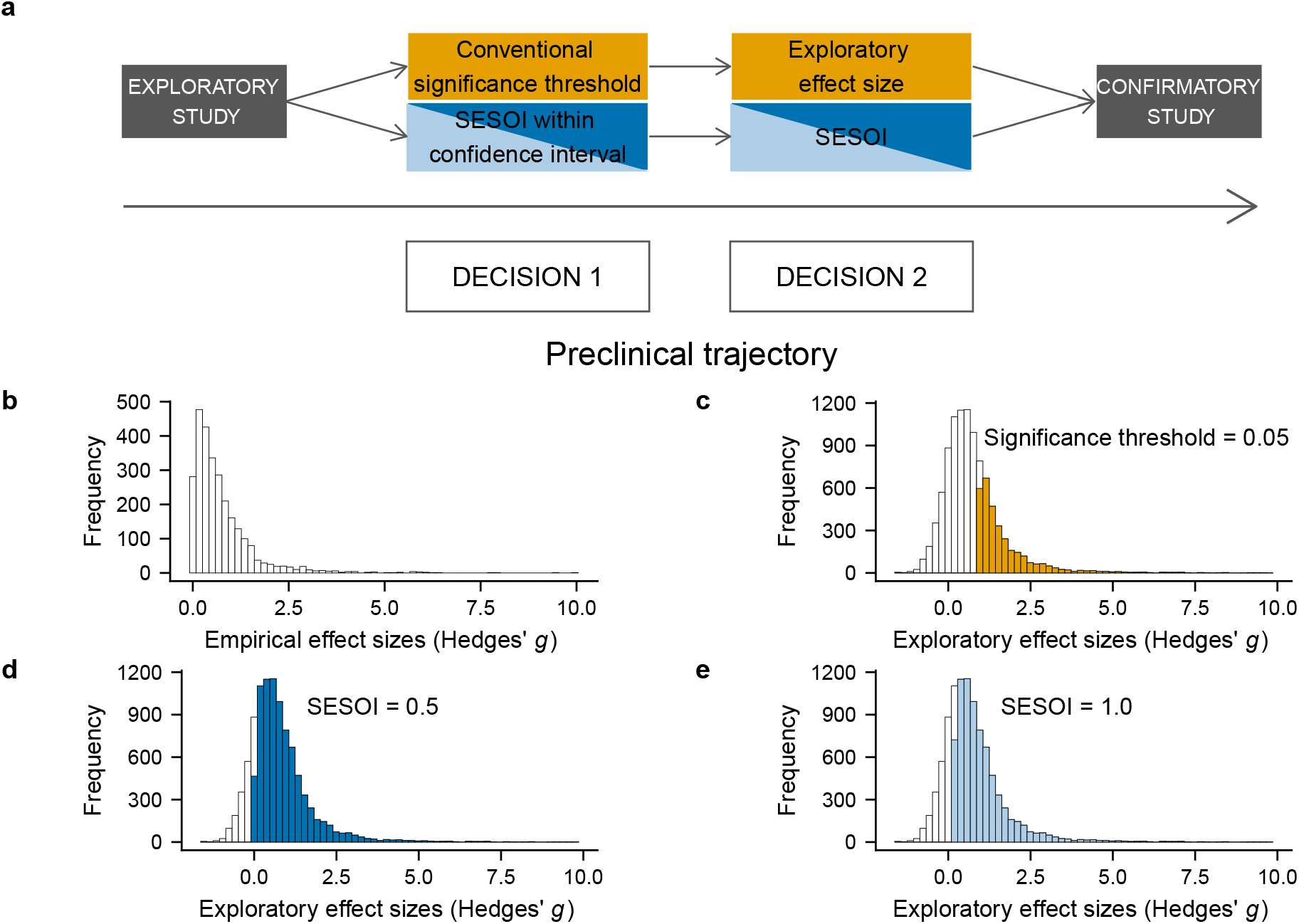
Preclinical research trajectory and transition rates. **a**, Along the trajectory, two decisions have to be made. DECISION 1: Which experiments should move from exploration to confirmation? DECISION 2: How should the sample size for a confirmatory study (i.e. within-lab replication) be estimated? The yellow and blue panels display the decision criteria and approaches to estimate the sample size along the two tajectories that were compared in this study (yellow = Standard; blue = SESOI). **b**, Distribution of empirical effect sizes (*n* = 2729) extracted from Bonapersona *et al*., 2021. **c-e**, Exploratory effect sizes (*n* = 9958, values > 10 were removed for better display). The shaded bars show the effect sizes that were detected using one of the two decision criteria. **c**, Yellow shaded bars indicate those effect sizes that were identified for confirmation using the conventional significance threshold (*α* = .05). **d**, Dark blue shaded bars indicate effect sizes that were selected using a SESOI of 0.5. **e**, Light blue shaded bars indicate effect sizes that were selected using a SESOI of 1.0.

We compared the two trajectories with regard to how well they meet the complementary goals of exploration and confirmation. Specifically, we were interested in the number of studies that transition from exploration to confirmation, the number of animals needed in confirmation as well as the false negative rate (FNR), false positive rate (FPR), positive predictive value (PPV), and negative predictive value (NPV).

## Results

### Transition rates from exploration to confirmation

Of the 10000 simulated exploratory studies, more experiments were selected for transition to the confirmatory phase in the SESOI trajectory compared to the Standard trajectory. Thus, in the SESOI trajectory 83% (SESOI = 0.5) and 75% (SESOI = 1.0) of experiments transitioned to confirmation whereas 33% of experiments proceeded in the Standard trajectory (Fig. 1c–e). Importantly, we compared how reliably each of the two trajectories detected effects of a given size. As reference effect sizes, we chose 0.5 and 1.0. to be our effect sizes of interest. In the underlying true effect size distribution that we used for our simulation, 50% were *≥* 0.5 and 24% were *≥* 1.0. Of those, 97% and 99% transitioned to confirmatory stage using a SESOI of 0.5 and 1.0, respectively. Using a significance threshold of *α* = .05 resulted in 57% and 84% correctly identified effects. The transition rates demonstrate that the significance threshold risks discarding potentially meaningful effects at early stages of discovery particularly if effects of interest are below 1.0 (Fig. 2a). The SESOI criterion reduced false negatives compared to the conventional significance threshold, but at the same time made many more confirmatory studies necessary.

**Figure 2:**
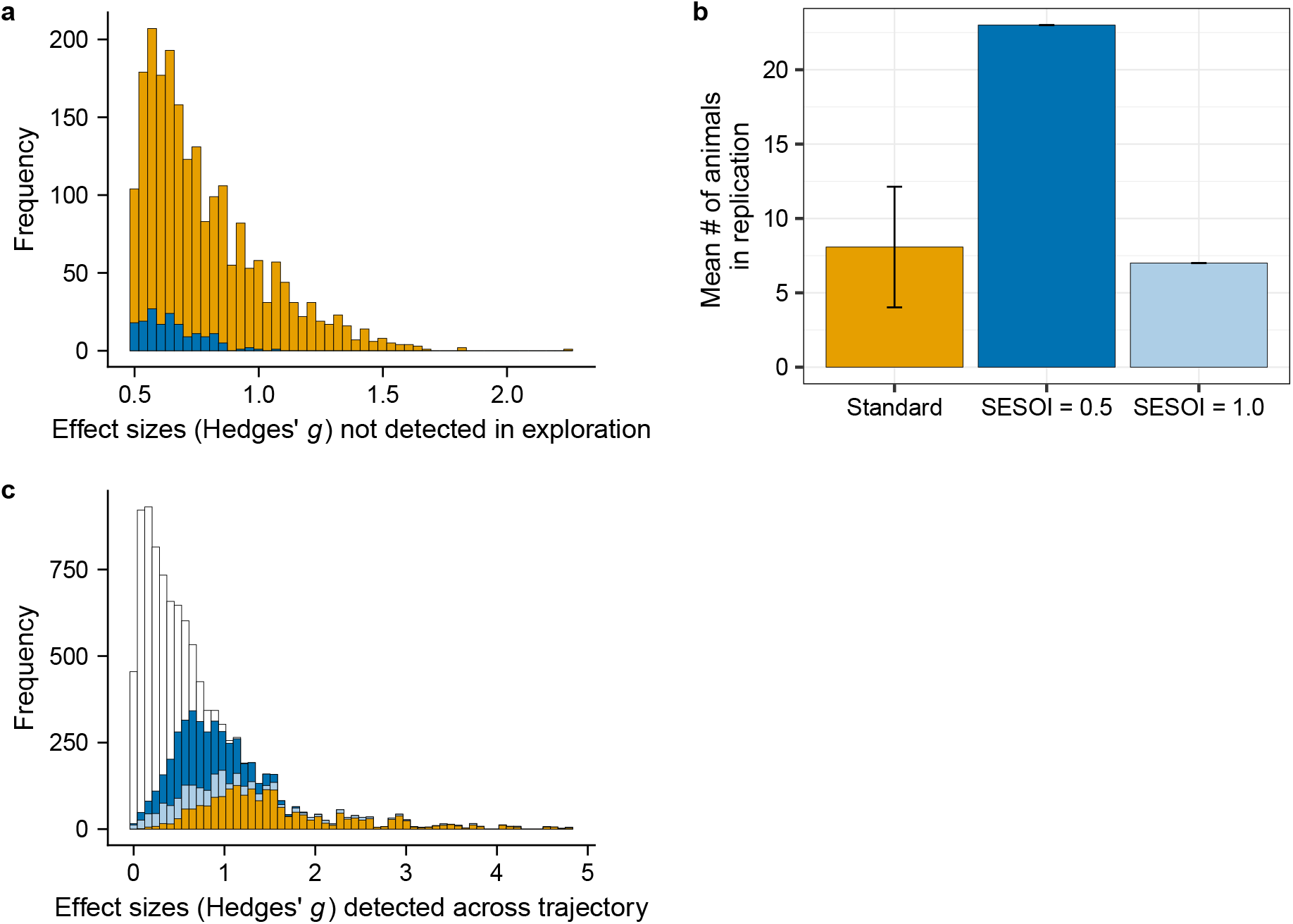
Sample sizes in confirmation and significant effect sizes after replication. **a**, Underlying effect sizes that were not detected during exploration using either the conventional significance threshold (*α* = .05, yellow shaded bars) or a SESOI of 0.5 (dark blue shaded bars). 2154 effects > 0.5 were falsely eliminated when the significance criterion was employed compared to 172 if a SESOI of 0.5 was applied. **b**, Number of animals needed in the confirmatory study. In the Standard trajectory, sample sizes are low (*n*_*rep*_ = 8.08 *±* 4.05), as they are based on large exploratory effect sizes. Error bars represent standard deviations. In case of trajectories using a SESOI, the number of animals is fixed. Using a SESOI of 0.5 results in *n*_*rep*_ = 23, whereas a SESOI of 1.0 achieves a sample size comparable to those in the Standard trajectory (*n*_*rep*_ = 7). All sample sizes displayed and reported in the text are the number of animals needed in *each* group (control and intervention). **c**, The histogram displays the distribution of sampled effect sizes (*n* = 9875, values *>* 5 were removed for better display), that constituted our underlying true effect sizes in the simulation. Shaded bars indicate which underlying effect sizes were detected in the two-stage process. Dark blue shaded bars represent effect sizes that were detected across the SESOI trajectory employing a SESOI of 0.5. Light blue bars represent effect sizes that were detected employing a SESOI of 1.0. Yellow shaded bars indicate effect sizes that were detected across the Standard trajectory. As the histogram illustrates, using the Standard trajectory for discovery is not efficient as it overlooks numerous potentially meaningful effects.

### Sample size calculation for confirmation

In a subsequent step, we estimated the sample size for a confirmatory study. In the Standard trajectory this was done via a power analysis using the initial exploratory effect size. The SESOI trajectory used the pre-defined SESOI that was employed as decision criterion earlier. Importantly, we set the power to detect an effect of the size of our SESOI to 50% (for more details see Methods section).

In the Standard trajectory, this resulted in small sample sizes per group (Standard: *n*_*rep*_ (*M ± SD*) = 8.08 *±* 4.05; Fig. 2b). In the SESOI trajectory, the number of animals varied with the SESOI that was chosen (SESOI 0.5: *n*_*rep*_ = 23, SESOI 1.0: *n*_*rep*_ = 7; Fig. 2b). The small numbers in the Standard trajectory reflect the large exploratory effect sizes that passed on to confirmation and formed the basis for confirmatory sample size estimation (Fig. 1c).

### Outcomes across trajectory

To estimate discovery success, we calculated the positive predictive value (PPV), negative predictive value (NPV), false positive rate (FPR), and false negative rate (FNR) across both trajectories. The PPV of a study is the post-study probability that a positive finding which is based on statistical significance reflects a true effect.^10^ Likewise, the negative predictive value (NPV) is the post-study probability that a negative finding reflects a null effect.^8^ Both PPV and NPV are calculated from the prior probability (or prevalence), as well as the sensitivity and specificity of the test. In our study, the prior probability of an effect of a given size (Hedges’ *g* of 0.5 and 1.0, respectively) was calculated from the empirical effect size distribution. For example, to calculate the prior probability of an effect Hedges’ *g* = 0.5, we divided the number of effects *≥* 0.5 by the total number of effects in the distribution. For an effect Hedges’ *g* = 0.5, the prior probability was 0.5, for Hedges’ *g* = 1.0, the prior probability was 0.24.

As Fig. 2c and Fig. 3 illustrate, the two trajectories exhibit distinct strengths. The Standard trajectory successfully weeds out effects smaller than 0.5 and 1.0 but detects less than half and two thirds of the relevant effect sizes, respectively. Vice versa for the SESOI trajectory which successfully identifies relevant effect sizes but is less specific. The PPV reaches higher levels across the Standard trajectory (Standard 0.5: 0.97, Standard 1.0: 0.76) compared to the SESOI trajectory (SESOI 0.5: 0.84, SESOI 1.0: 0.62; Fig. 3a). This means that among the significant results after the two-stage process a higher proportion reflects a true effect in the Standard trajectory. The NPV favored the SESOI trajectory over the Standard trajectory, particularly for lower SESOI (SESOI 0.5: 0.83, SESOI 1.0: 0.93; Standard 0.5: 0.62, Standard 1.0: 0.9; Fig. 3b). The FPR was consistently higher in the SESOI trajectory compared to the Standard trajectory (SESOI 0.5: 0.16, SESOI 1.0: 0.15, Standard 0.5: 0.01, Standard 1.0: 0.06; Fig. 3c). The FNR revealed the most pronounced differences between the trajectories. It was considerably lower in the SESOI trajectory for smaller SESOI (SESOI 0.5: 0.17, SESOI 1.0: 0.21, Standard 0.5: 0.61, Standard 1.0: 0.34; Fig. 3d).

**Figure 3:**
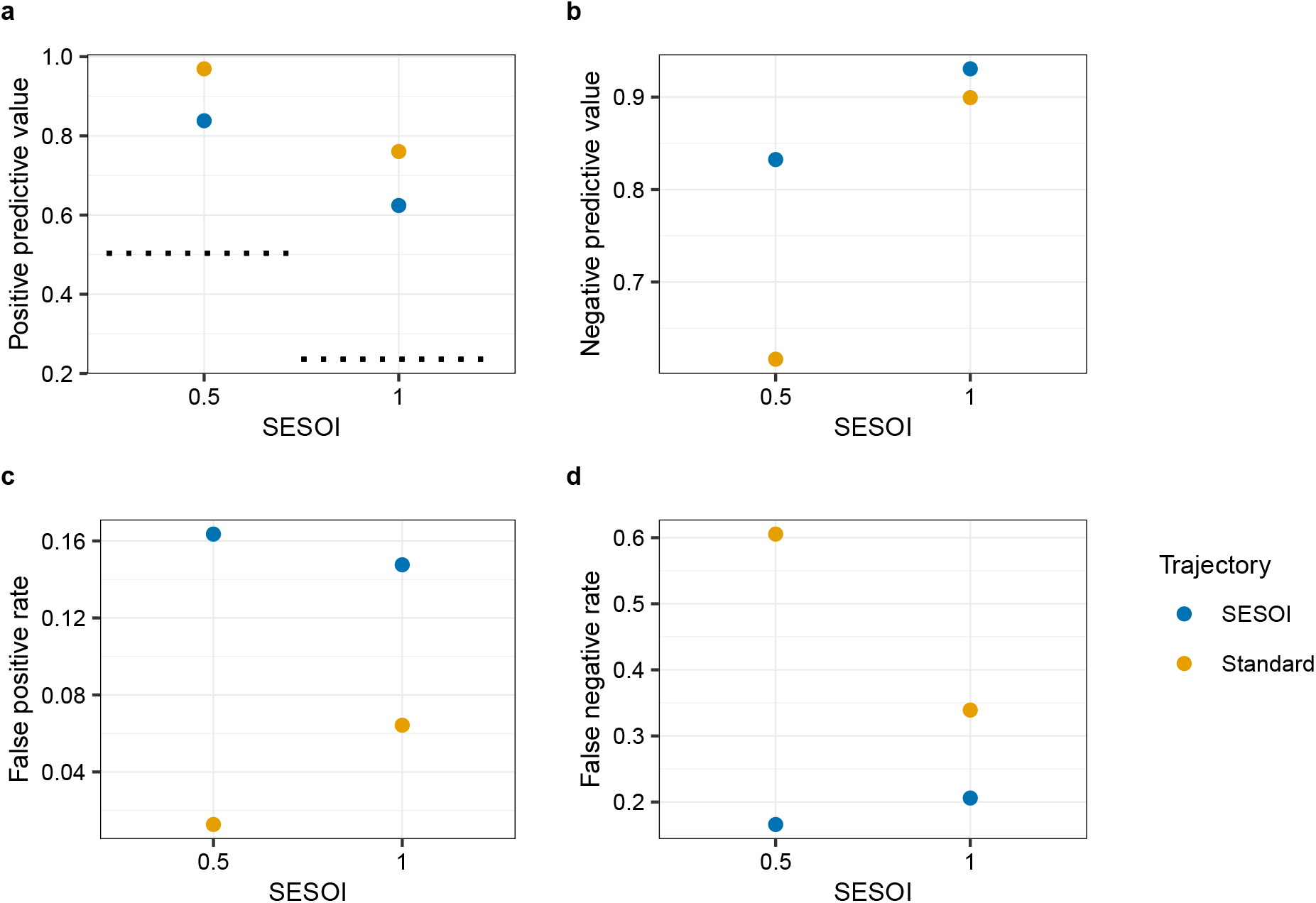
Outcomes across trajectory. **a**, The positive predictive value across the trajectory is consistently higher in the Standard trajectory. **b**, The negative predictive value is higher in the SESOI trajectory, but the difference between the two trajectories decreases for larger SESOI. **c**, The false positive rate across the trajectory is better controlled in the Standard trajectory. **d**, The false negative rate across the trajectory is much higher in the Standard trajectory but decreases as SESOI increases.

### Resource reallocation from SESOI to Standard trajectory

The results regarding the FNR across the trajectory raised the question whether we would be able to increase sensitivity in the Standard trajectory if we reallocated the resources used in the SESOI trajectory. Specifically, if we took the total amount of experimental units (EU) used in the SESOI trajectory (given a SESOI of 0.5 this would amount to 10 EU * 2 [groups] * 10000 [studies] in exploration + 23 EU * 2 [groups] * 8333 [studies] in confirmation), we could run 10000 exploratory studies with 30 EU in *each* group in the Standard trajectory. How would an increased sample size at this early stage change the PPV, NPV, FPR, and FNR? We compared outcomes across the two-stage SESOI and Standard trajectories with outcomes of the 30 EU exploratory study in Standard trajectory. As displayed in Fig. 4a, an increase in initial sample size benefits the sensitivity indicated by a sharp drop in false negatives (FNR: 0.13) and an elevated NPV (NPV: 0.86). However, this comes at the cost of a higher FPR (FPR: 0.18) and a decreased PPV (PPV: 0.83).

**Figure 4:**
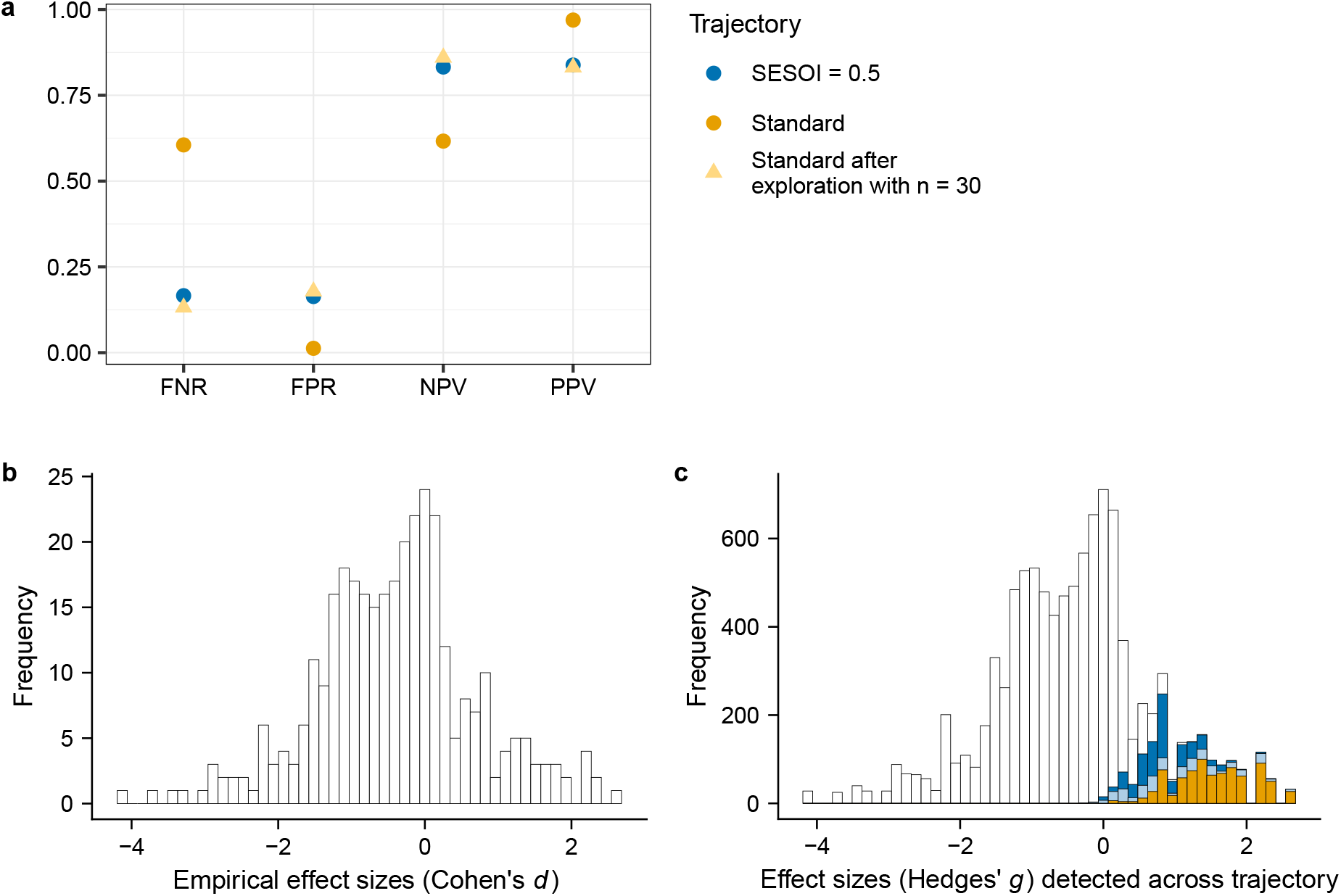
Outcomes of resource reallocation and significant effect sizes after replication for pessimistic effect size distribution. **a**, Increasing initial sample size in the Standard trajectory benefits sensitivity indicated by a lower FNR. Overall, the results are comparable with the two-stage SESOI trajectory using a SESOI of 0.5. **b**, Distribution of empirical effect sizes (*n* = 336) extracted from Carneiro *et al*., 2018. Here, the standardized mean difference was expressed as Cohens *d*. **c**, The histogram displays the distribution of sampled effect sizes (*n* = 10000), that constituted our underlying true effect sizes in the simulation. Shaded bars indicate which underlying effect sizes were detected in the two-stage process (SESOI = 0.5: dark blue shaded bars, SESOI = 1.0: light blue shaded bars, Standard: yellow shaded bars). As the histogram illustrates, the Standard trajectory does not successfully identify relevant effects.

### Pessimistic effect size distribution

In addition to the simulation based on the effect size distribution by Bonapersona *et al*., (2021; in the following referred to as optimistic distribution),^9^ we ran the simulation also with a more pessimistic published effect size distribution^11^ (Fig. 4b). The main difference is that this distribution contains negative effect sizes and that the prior probabilities for an effect of 0.5 and 1.0 are considerably lower (Hedges’ *g* = 0.5: 0.17, Hedges’ *g* = 1.0: 0.1). The SESOI trajectory is more sensitive and successfully identifies relevant effect sizes compared to the Standard trajectory (Fig. 4c). The results regarding the PPV and NPV are very similar to those obtained based on the optimistic distribution (PPV: SESOI 0.5: 0.89, SESOI 1.0: 0.73, Standard 0.5: 0.98, Standard 1.0: 0.82; NPV: SESOI 0.5: 0.98, SESOI 1.0: 0.98, Standard 0.5: 0.9, Standard 1.0: 0.96; Fig. 5a-b). However, in the pessimistic scenario, the SESOI trajectory performs considerably better with regard to the FPR (SESOI 0.5: 0.02, SESOI 1.0: 0.03; Fig. 5c) although the FPR is still slightly lower in the Standard trajectory (Standard 0.5: 0.002, Standard 1.0: 0.02). The FNR remains concerningly high in the Standard trajectory (Standard 0.5: 0.53, Standard 1.0: 0.32; Fig. 5d), whereas in the SESOI trajectory it is decreased compared to the optimistic distribution (SESOI 0.5: 0.12, SESOI 1.0: 0.16). Transition rates and sample sizes in confirmation are displayed in Fig. S12.

**Figure 5:**
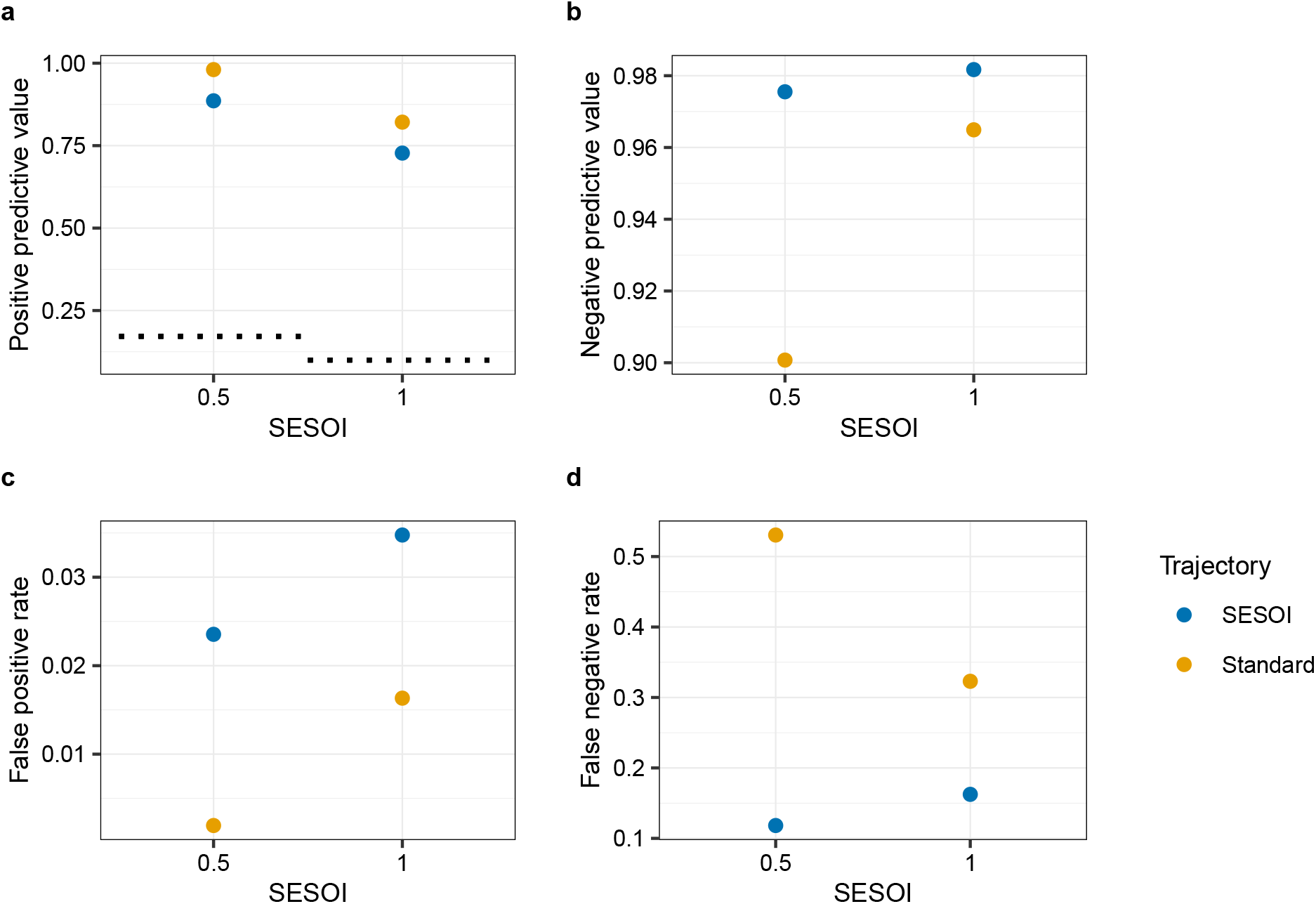
Outcomes across trajectory based on pessimistic effect size distribution. **a**, The positive predictive value across the trajectory is consistently higher in the Standard trajectory. **b**, The negative predictive value is higher in the SESOI trajectory. **c**, The false positive rate across the trajectory is well controlled in both trajectories. **d**, The false negative rate is much lower in the SESOI trajectory. Across the Standard trajectory, more than half (SESOI = 0.5) and one third (SESOI = 1.0) of the relevant effect sizes are missed, respectively.

## Discussion

Overall, our simulation shows that current practice reflected by the Standard trajectory does not meet the complementary goals of exploration and confirmation along a preclinical research trajectory. In the Standard trajectory, the switch from exploratory to confirmatory mode eliminates numerous potentially meaningful effects. A more lenient significance threshold (*α* = 0.1) elevated the number of studies that proceeded to confirmation (Fig. S10). However, not to an extent that appreciably impacted the PPV and NPV (Fig. S11). The same holds for a slight decrease/increase of the initial sample size (Fig. S4–S5). When we tested a more pessimistic published effect size distribution results were preserved (Fig. 5). Overall, the decision criterion used to decide whether to transition to confirmation has been shown to have a stronger impact on our outcomes of interest than the approach for confirmatory sample size estimation (Fig. S9). In light of our findings, we advise a reconsideration of planning and conducting experiments in early-stage preclinical research.

Whereas the Standard trajectory excels in specificity as indicated by a low FPR and reflected by a high PPV, its low sensitivity should cause concern. Upon exploration, a considerable number of potentially meaningful effects is eliminated (Fig. 2a). Unlike in our simulation, in real life we cannot know how many meaningful effects were falsely discarded. The costs of false negatives are therefore difficult to quantify. This might explain why preventing false positives is often a greater concern for researchers. However, as Kane and Kimmelman, (2021)^8^ point out, false negatives are just as important to consider. The focus needs to shift towards more sensitive screening strategies to prevent false negatives, especially in research areas that have yielded few advances with regard to effective treatments.^3,12–16^

Low sensitivity in the Standard trajectory could be remedied by using more animals in the exploratory study (Fig. 4a). However, that might undermine the tentative nature of exploration. Furthermore, conducting an additional confirmatory study (as in the two-stage SESOI trajectory) does not only allow to increase reliability but also enables a gradual refinement of experimental design using measures like randomization and blinding to reduce bias.^5,17^

Our results underscore the need for a systematic estimation of true effects using sensitive screening strategies. We therefore propose to proceed following these steps when embarking on an investigation: 1. Choose a SESOI. The rationale for the choice of a SESOI should be transparently reported and can be based on various considerations, like feasibility given practical constraints or meta-analytic effect size distributions in your field.^18^ 2. Conduct an exploratory study and apply the pre-specified SESOI as decision criterion to decide whether to move forward to confirmation. Be aware that the width of your CI is a function of the sample size, i.e., the less animals you use, the wider the CI. The CI, in which the SESOI should fall, needs potentially be adjusted based on the reliability of the initial study (Fig. S1). This has to be done also *a priori*. 3. If the decision points to a transition to confirmatory mode, use the SESOI to calculate the confirmatory sample size with a power of .50, ideally using a one-sided test. 4. Conduct a confirmatory study. Here, we chose significance as a measure of confirmation success. However, other methods such as the estimation of effect sizes and their CI or the sceptical *p*-value^19^ might be more informative.

Our results are dependent on underlying assumptions represented by the empirical effect size distributions from Bonapersona *et al*. (2021) and Carneiro *et al*. (2018). If effect sizes in a field considerably deviate from these distributions (i.e., are larger), following our recommendation might be dispensable, as the issue of low sensitivity to detect effects does not arise. Our method specifically addresses preclinical research areas that need to weigh the number of animals against the likelihood of inference errors. This is the case in fields where effect estimates are expected to be small. This applies for example if a new treatment is tested against a (standard of care) comparator.

Apart from the outcomes, using a SESOI for study planning comes with a crucial advantage: It requires researchers to explicitly state and motivate which effects they consider relevant. If we took the conceptual implications of null hypothesis significance testing literally, every effect > 0 would be deemed meaningful. In reality, we believe, this is not the case. Researchers always need to balance their available resources against the potential benefit of an experiment. This process involves deliberations about which effect sizes are meaningful and worthwhile to confirm. However, only rarely are these considerations transparently stated. We suppose that the explicit choice of a SESOI fosters conceptual deliberations and thereby improves experimental design. In addition, a clear conception of a meaningful effect might be of benefit in the communication with regulatory authorities.

We are exploring solutions to a complex problem. We acknowledge that the solution cannot be simple and that there is no one-size-fits-all. The two trajectories compared here have different strengths and weaknesses. Thus, they might be used under different circumstances. We concede that applying a SESOI is more costly, yet our simulation shows that statistical significance even with a relaxed threshold is not a sensible criterion to screen for potentially meaningful effects during exploration. Sensitivity is an indispensable requirement for successful discovery. Thus, in fields that have made little progress with regard to effective treatments, sensitivity should be prioritized over specificity to eventually generate patient benefit.

The method we present here is an easily applicable alternative to current practice in preclinical animal research. Optimizing decision criteria by employing a SESOI increases chances to detect true effects while keeping the number of animals at a minimum.

## Methods

### Simulation

We explored different approaches to perform preclinical animal experiments via simulations. To this end, we modeled a simplified preclinical research trajectory from the exploratory stage to the results of a confirmatory study (within-lab replication; Fig. 1a). Along the trajectory, there are different ways to increase the probability of not missing potentially meaningful effects. After an initial exploratory study, a first decision identifies experiments for replication. In our simulation, we employed two different decision criteria that indicate when one should move from the exploratory to confirmatory mode. If a decision has been made to replicate an initial study, we applied two approaches to determine the sample size for a replication study (smallest effect size of interest (SESOI) and power analysis based on the exploratory effect size), as outlined in detail below.

#### Empirical effect size distribution

Simulations were based on an empirical effect size distribution from the recently published literature.^9^ From this distribution, we were able to calculate the prior probability of a particular effect. For example, if we wanted to know the prior probability of an effect Hedges’ *g* = 0.5, we divided the number of effects *≥* 0.5 by the total number of effects in the distribution.

The distribution of effect sizes extracted from Bonapersona *et al*., (2021) contains 2738 effect sizes from the fields neuroscience and metabolism. All effect sizes are calculated as the standardized difference in means (Hedges’ *g*). Effect size estimates range from 0 to 24.61 and have a median of 0.5.

As a robustness check, we ran the simulation also with a more pessimistic published effect size distribution.^11^ The study by Carneiro *et al*., (2018) systematically examined effect sizes in the rodent fear conditioning literature. The distribution consists of 336 effect sizes calculated as the standardized mean difference (Cohen’s’ *d*). The effect sizes range from -4.14 to 2.6, with a median of -0.38. In our simulation, standardized mean differences were calculated as Hedges’ *g*.

#### Exploratory mode

From the empirical effect size distribution, we drew 10000 samples of effect sizes from which we created 10000 study data sets. Each data set comprised data of two groups consisting of ten experimental units (EU) each drawn from a normal distribution. We chose a number of ten EUs based on reported sample sizes in preclinical studies.^9,20^ Our simulated design mimics a comparison between two groups where one group receives an intervention and the other serves as a control group. The difference between the intervention and control group in each study data set was determined by the effect size that was drawn from the empirical effect size distribution. In that way, we ensured that our simulated data was comparable to data in the published literature in the respective fields (neuroscience and metabolism). In the simulated exploratory study, the study data sets are compared using a two-sided two-sample Welch’s *t*-test. From these exploratory study results, we extracted *p*-values, exploratory effect size estimates, and their 95% confidence intervals (CI). We then employed two different criteria based on the *p*-value or 95% CI, respectively, to decide whether to continue to confirmatory mode.

#### Decision criteria to proceed to confirmation

The first decision criterion employed the conventional significance threshold (*α* = .05) to decide whether to replicate an exploratory study. If a *p*-value extracted from a two-sided two-sample Welch’s *t*-test was *≤* .05 and the exploratory effect size was > 0, this study proceeded to confirmation. If not, the trajectory was terminated after the exploratory study. We chose this decision criterion as our reference, as this is what we consider to be current practice.

As an alternative to this approach, we propose to set a smallest effect size of interest (SESOI) and examine whether the 95% CI around the exploratory effect size estimate covers this SESOI. A SESOI is the effect size that the researcher based their domain knowledge and given practical constraints considers biologically and clinically meaningful.^18^ Our proposed method is similar to a non-inferiority test. In our case the null hypothesis stated that the exploratory study identified an effect that is inferior to our pre-defined SESOI. If this was not the case for an exploratory study, i.e., if the CI around the exploratory effect size covered our SESOI, non-inferiority was established, and the study was carried forward to confirmatory mode. Again, exploratory effect sizes < 0 were not considered for confirmation. Importantly, we did not consider significance in this approach. In our simulation, we used Hedges’ *g* = 0.5 and Hedges’ *g* = 1.0 as SESOI. This approach emphasizes the importance of effect sizes rather than statistical significance to evaluate an intervention’s effect. Further, we expected this approach to be more lenient than statistical significance and to allow a broader range of effect sizes to pass on to be further investigated.

Both decision criteria identified exploratory effect sizes < 0 to transition to confirmation. This would mean that an experiment which showed an effect in favor of the control would be replicated. As we think this is an unrealistic scenario, negative effect sizes were not carried forward to confirmation in the simulation. Based on the underlying effect size, these negatives were categorized as false negatives or true negatives to later calculate the outcomes across the two trajectories.

#### Approaches to determine sample size for replication

Once the decision to continue to confirmatory mode has been made, we employed two different approaches to determine the sample size for the confirmatory study. After the exploratory study, only effect sizes that showed an effect in favor of the treatment were considered for further investigation. Thus, for the confirmatory study, a one-sided two-sample Welch’s *t*-test was performed. In the Standard trajectory, the desired power level for the confirmatory study was set to 80%, *α* was set to .05. To determine the sample size given power and *α*, we used the exploratory effect size estimate. In the SESOI trajectory, we employed the same SESOI used as decision criterion earlier to calculate the confirmatory sample size. Our SESOI was set such that the confirmatory study has a power of 50% to detect an effect of this size. This power level was chosen to ensure that the likelihood of a false positive finding below the threshold determined by our SESOI is negligible. The aim during confirmation is to weed out false positives. Additionally, 0.5 is only the *smallest* effect size we are interested in, meaning all other effect sizes of interest (i.e., effect sizes > 0.5) have a higher chance of being detected.^21^ This increase in power with increasing effect size is larger with a power level of 50% compared to a power level of 80% as illustrated by the power curves in Fig. S2.

#### Confirmatory mode

For each of the studies that met the decision criterion after the exploratory study (either *p ≤* .05 or SESOI within the 95 % CI of the exploratory effect size estimate), a confirmatory study was performed. The number of studies conducted varied with the decision criterion used and, in case of the criterion employing a SESOI, also with the SESOI (0.5 and 1.0). A confirmatory study was performed as a one-sided two-sample Welch’s *t*-test, where the number of animals in each group was determined by the approach to determine the sample size. For a confirmatory study to be considered “successful”, the *p*-value had to be below the conventional significance threshold (*α* = .05).

#### Outcome variables

We compared the two trajectories (Standard and SESOI) regarding the transition rates from exploration to confirmation, number of animals needed in the confirmatory study, and positive predictive value (PPV), negative predictive value (NPV), false positive rate (FPR), and false negative rate (FNR) across the trajectories. Importantly, the PPV and NPV were calculated from the known prior probability (given by the empirical effect size distribution), the sensitivity (true positive rate), and the specificity (true negative rate). To make the rates comparable for the Standard and SESOI trajectory, we did not specify a non-existent effect as zero, but as an effect that is smaller than our SESOI (0.5 and 1.0, respectively). A true positive thus is a significant result that reflects an underlying effect of at least 0.5 (or 1.0, if our SESOI was 1.0). If a significant result represents an underlying effect smaller than 0.5 (or 1.0), this is categorized as a false positive (likewise for true and false negatives). This is a departure from the classical definition in the null hypothesis significance testing framework, where the benchmark against which is tested is zero.

#### Additional trajectories

In addition to the two trajectories compared in the main text, we also performed all simulations and analyses with the two crossed-over trajectories where each decision criterion is combined with the respective other approach to determine the sample size for confirmation. This resulted in the two trajectories SESOI-Standard (SESOI as decision criterion to move from exploration to confirmation + sample size estimation using the exploratory effect size estimate) and Standard-SESOI (Conventional significance threshold as decision criterion + SESOI for sample size estimation). Sample sizes in confirmation as well as outcomes across the trajectories are shown in Fig. S8 and Fig. S9.

#### Robustness checks

As robustness checks, we simulated all trajectories with an initial sample size (*n*_*init*_) of 5, 7 and 15 animals per group (in addition to 10 as described in the main part). We further performed all simulations and analyses with additional SESOI 0.1, 0.3, and 0.7. Outcomes are displayed in Fig. S4–S7. We also varied the significance threshold employed as decision criterion after exploration in the standard trajectory. The outcomes resulting from a more lenient significance threshold of *α* = 0.1 are displayed in Fig. S10 and Fig. S11. For the SESOI trajectory, we additionally varied the confidence interval around the exploratory effect size estimate used to determine whether to transition to confirmation. Transition rates and outcomes across trajectory are displayed in Fig. S1 and Fig. S2. Lastly, for the SESOI trajectory, we calculated the sample size for confirmation setting the power to .80 to investigate whether this would have beneficial effects on our outcomes of interest. Outcomes are shown in Fig. S3.

