## Supplementary figures for "Balancing sensitivity and specificity in preclinical research"

### Supplementary material to: Balancing sensitivity and specificity in preclinical research

#### Supplementary Figure 1

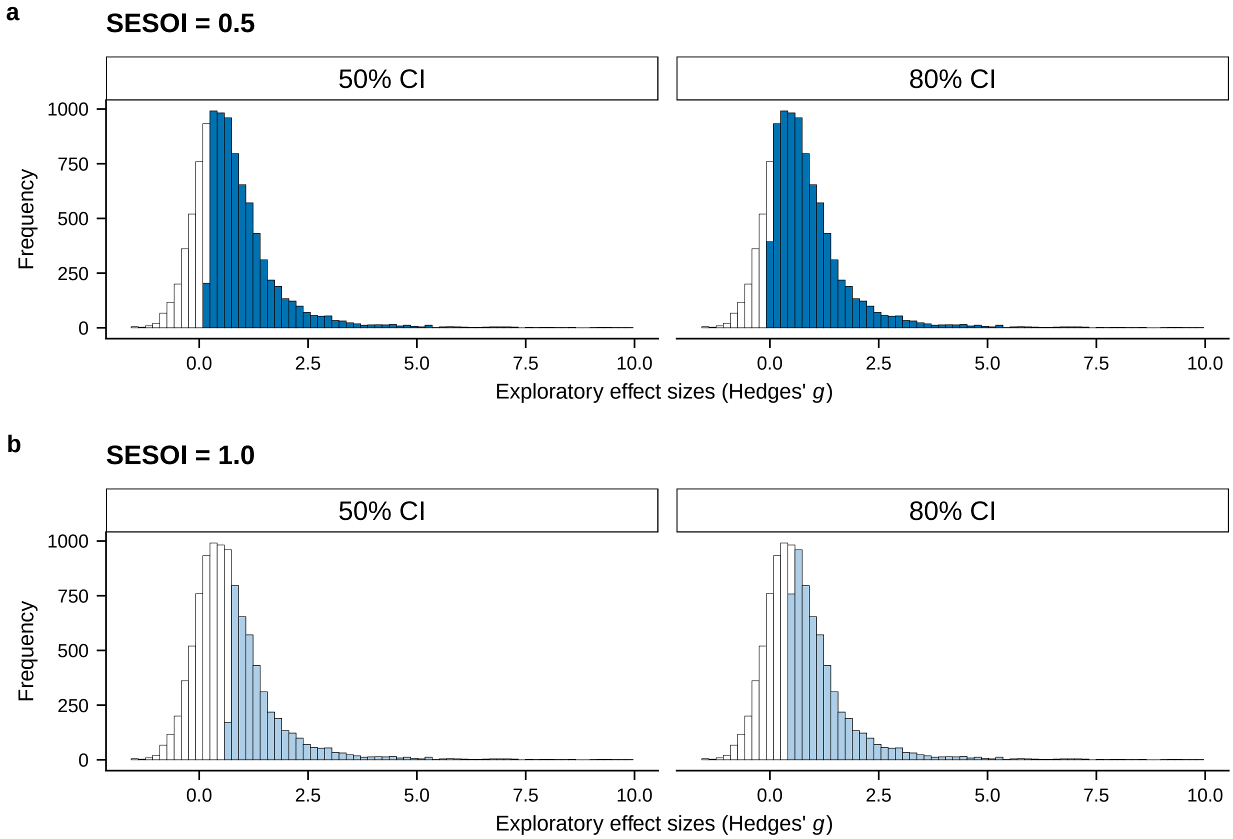


Figure S1: **Transition rates for two different confidence intervals and SESOI. a**, Given a SESOI of 0.5 and a 50% CI, 7209 effect sizes $> 0$ transition to the confirmatory stage. Employing an 80% CI increases the number to 8333 which is the same as for a 95% CI. **b**, Given a SESOI of 1.0 and a 50% CI, 4243 effect sizes $> 0$ transition to confirmation. Employing an 80% CI increases the number to 5790 which is still appreciably less than for a 95% CI. This showcases that depending on the SESOI used the bounds of the CI may have a crucial effect on the transition rate.

#### Supplementary Figure 2

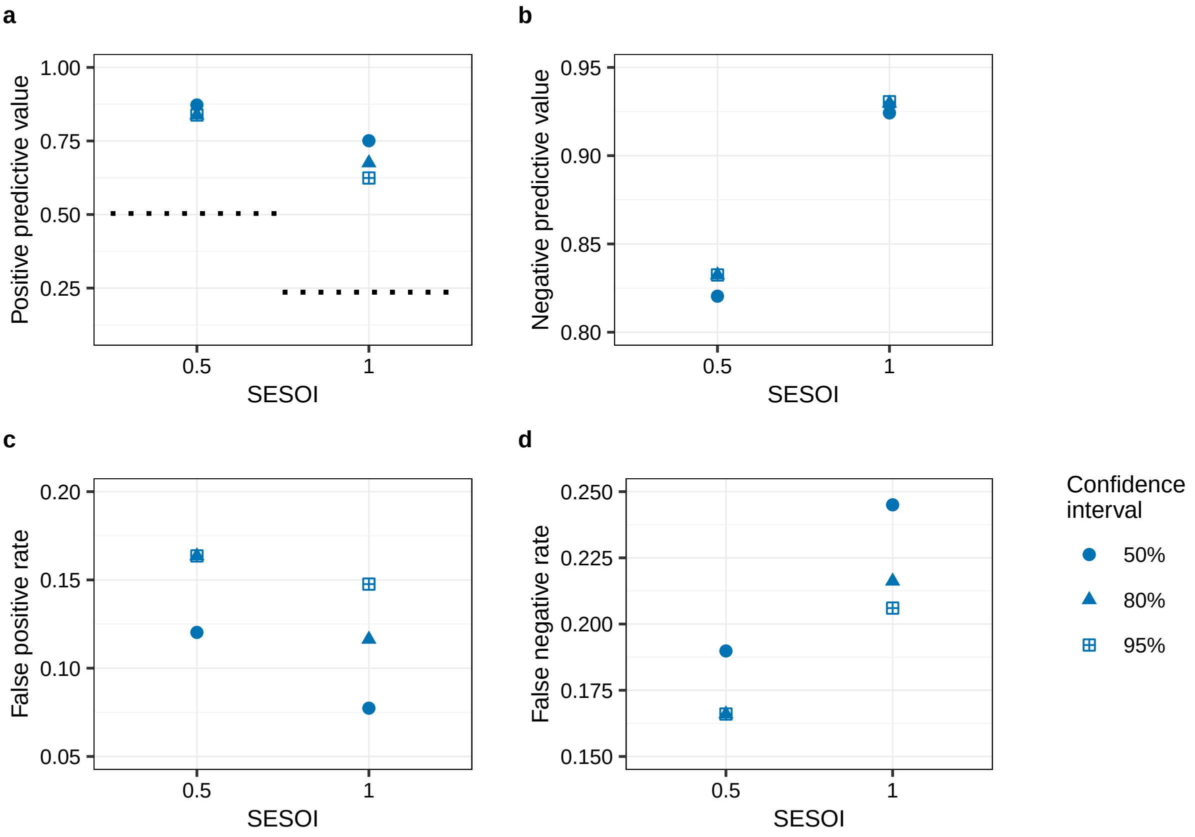


Figure S2: **Outcomes across SESOI trajectory for different confidence intervals and SESOI. a**, A narrower CI increases the PPV for larger SESOI. **b**, The NPV is slightly decreased if a 50% CI is employed. **c**, The FPR varies as a function of CI and SESOI. A narrower CI progressively reduces the FPR as SESOI increases. **d**, The FNR increases for narrower CI as SESOI increases.

#### Supplementary Figure 3

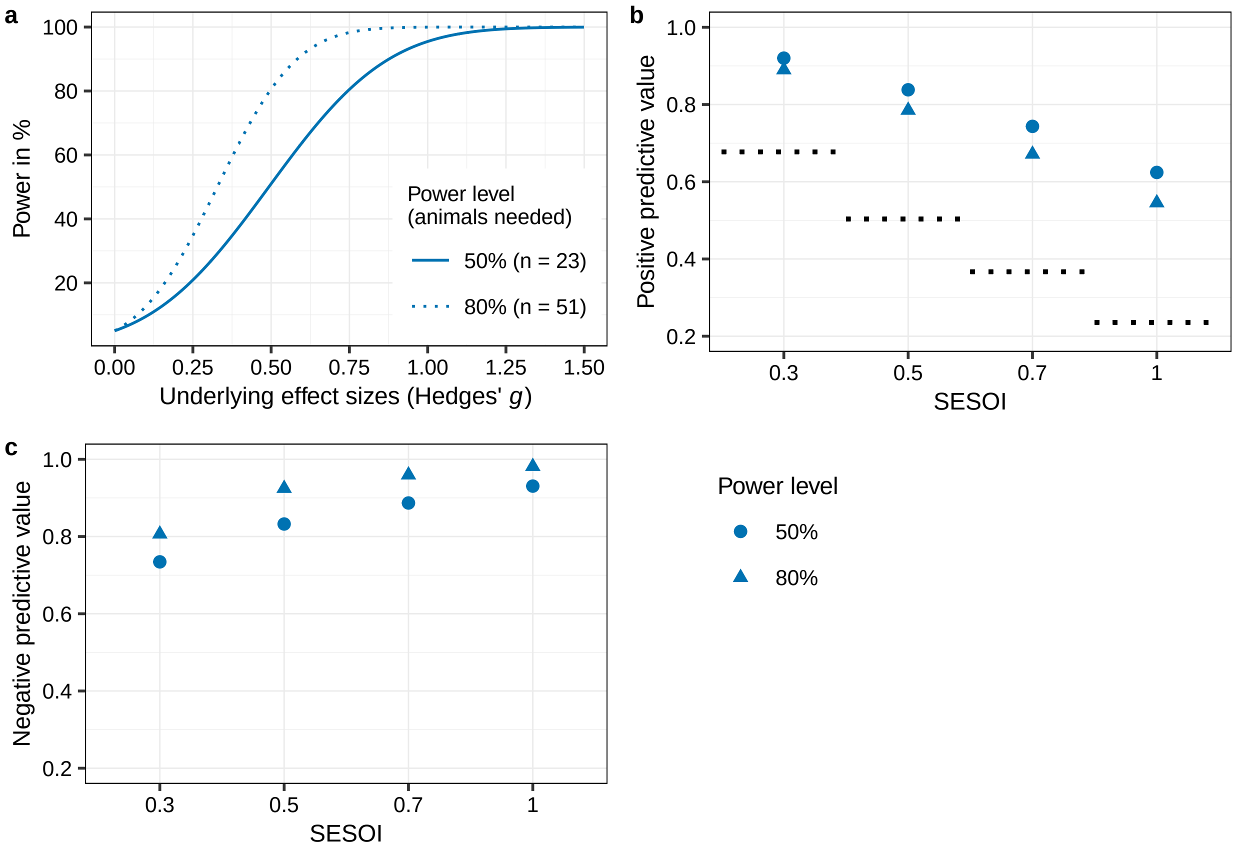


Figure S3: **Power curves, positive and negative predictive value across SESOI trajectory for two different power levels.** **a**, Power curves for two different power levels using a SESOI of 0.5 for sample size calculation. If the underlying effect size is 0.5, there is a 50% chance that this effect will be detected with a sample size of 23 animals per group and an 80% chance with 51 animals per group. If the underlying effect size is larger than the SESOI that was powered for, the slope of the 50% power curve (solid line) is steeper than the 80% curve. For example, for an effect size of 0.75, power increases by 30% (from 50% to 80%) whereas at a power level of 80% (dotted line) power increases by 18% (from 80% to 98%). However, in the latter case this increase comes at the cost of 28 more animals in *each* group. **b**, The PPV is slightly lower if the confirmatory study is powered at 80%. **c**, The NPV is consistently higher if study power is 80%.

#### Supplementary Figure 4

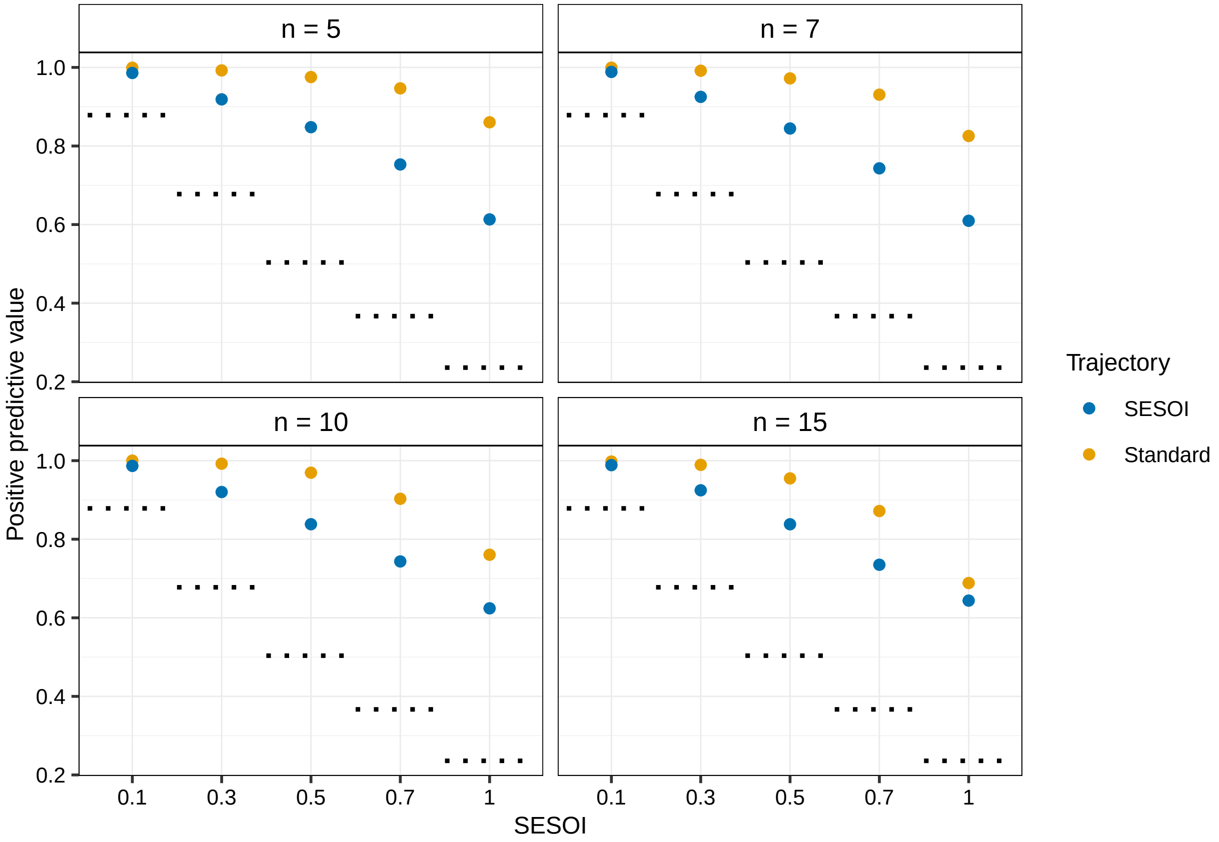


Figure S4: **Positive predictive value across trajectory for four different initial sample sizes.** The PPV is consistently higher in the Standard trajectory compared to the SESOI trajectory. An increase (or decrease) of the sample size in exploration does not change the direction of results. Whereas an increase of the initial sample size has very little impact in the SESOI trajectory, it seems to be detrimental in the Standard trajectory. Thus, for larger initial sample sizes the discrepancy between SESOI and Standard trajectory decreases.

#### Supplementary Figure 5

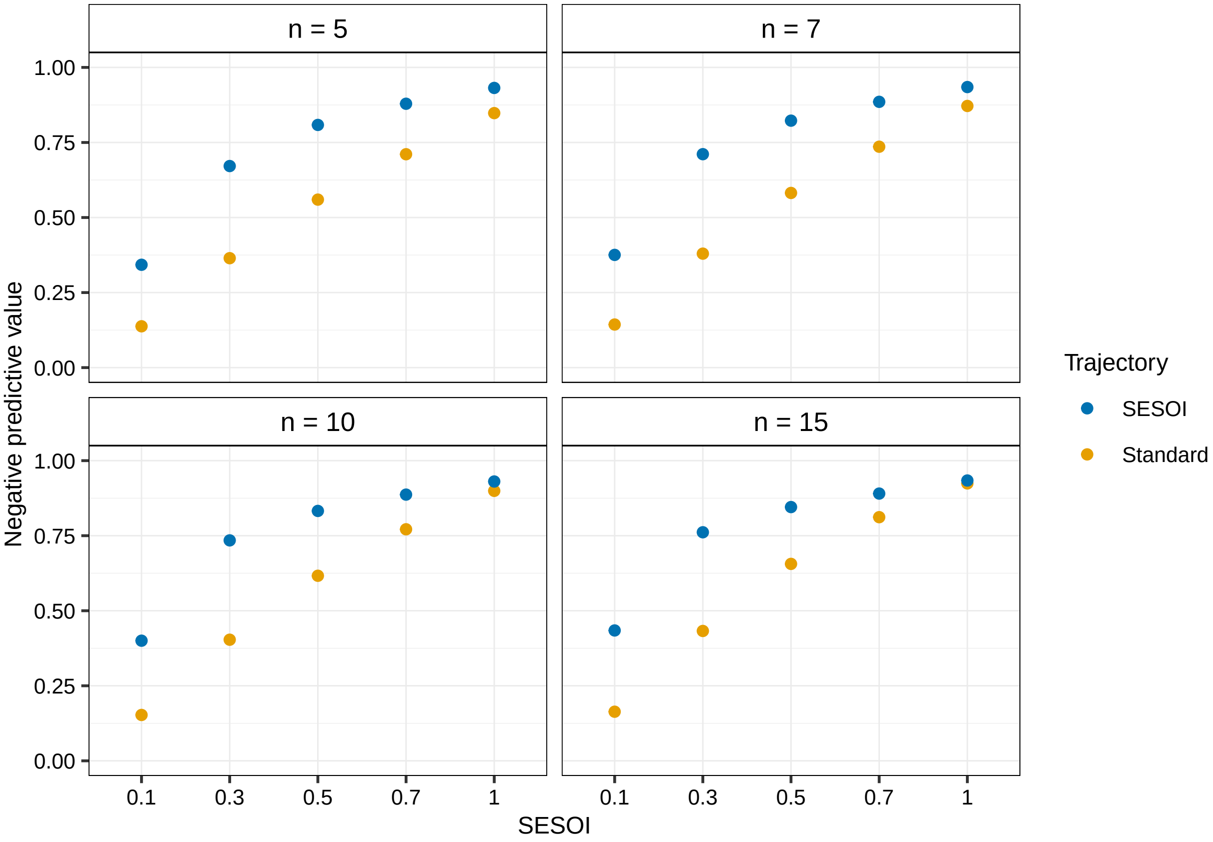


Figure S5: **Negative predictive value across trajectory for four different initial sample sizes.** The NPV is consistently higher in the SESOI trajectory compared to the Standard trajectory. An increase (or decrease) of the sample size in exploration does not change the direction of results. An increase of the initial sample size slightly increases NPV in the both trajectories. For larger initial sample sizes and SESOI the discrepancy between SESOI and Standard trajectory decreases.

#### Supplementary Figure 6

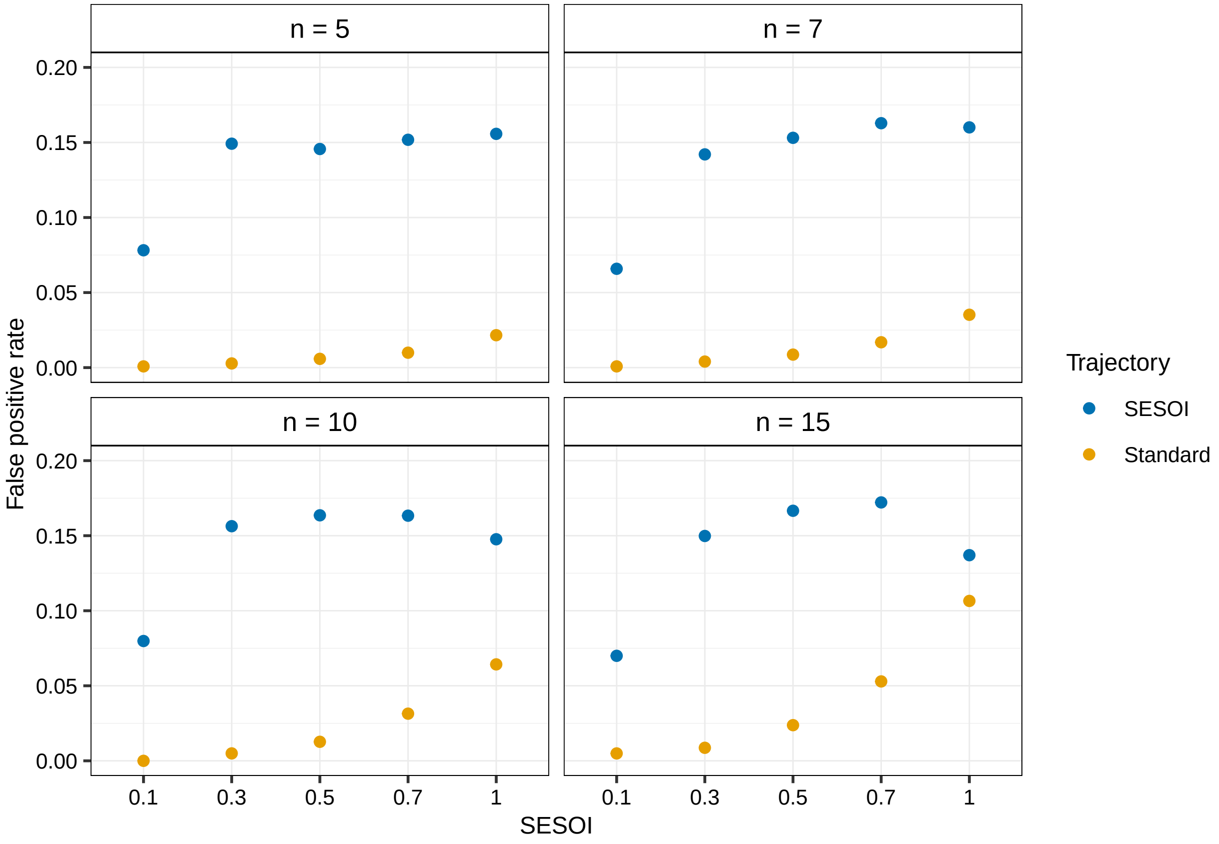


Figure S6: **False positive rate across trajectory for four different initial sample sizes.** The FPR is considerably higher in the SESOI trajectory across all SESOI. An increase of the initial sample size increases the FPR in the Standard trajectory, particularly for large SESOI.

#### Supplementary Figure 7

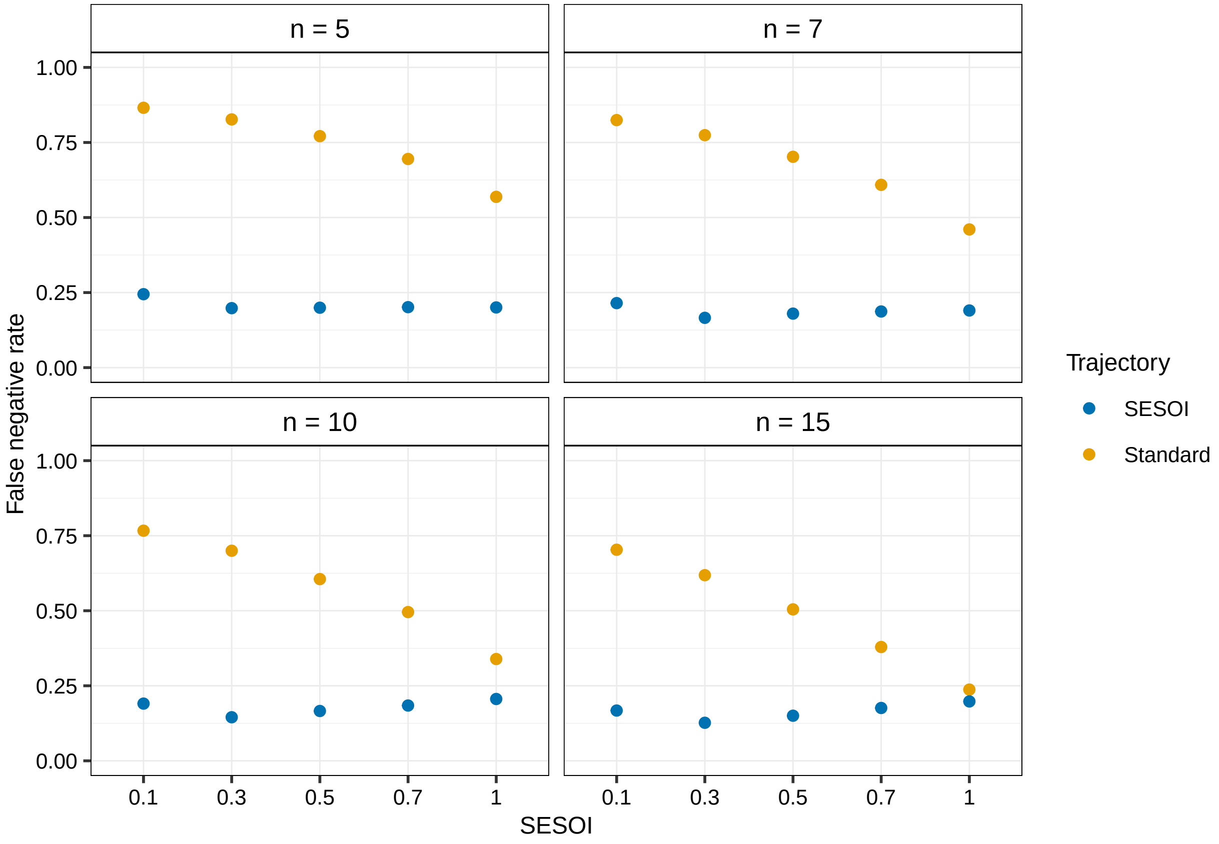


Figure S7: **False negative rate across trajectory for four different initial sample sizes.** Overall, the FNR in the SESOI trajectory is not strongly influenced by variation in initial sample size. However, an increase is beneficial in the Standard trajectory, particularly for large SESOI.

#### Supplementary Figure 8

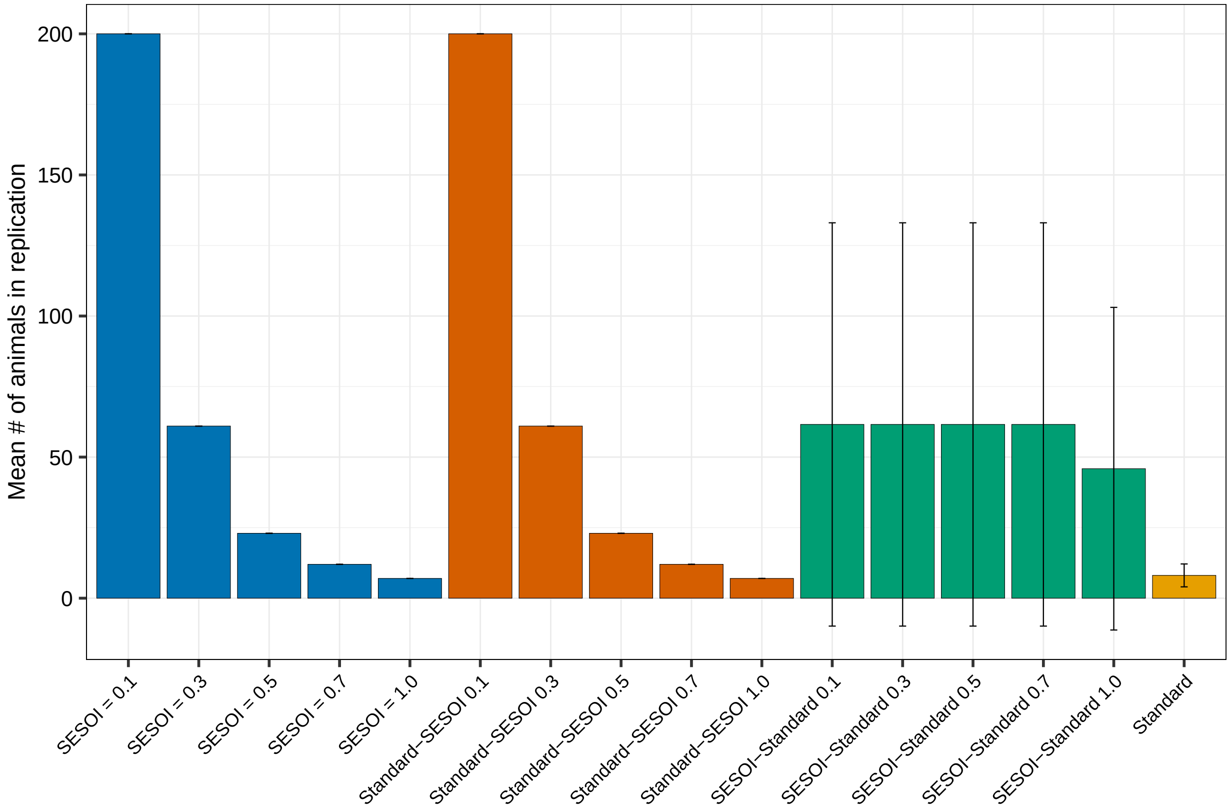


Figure S8: **Sample sizes in confirmation for four trajectories.** Sample sizes based on a SESOI are the same for SESOI and Standard-SESOI trajectory (SESOI 0.1: $n_{rep} = 200$, SESOI 0.3: $n_{rep} = 61$, SESOI 0.5: $n_{rep} = 23$, SESOI 0.7: $n_{rep} = 12$, SESOI 1.0: $n_{rep} = 7$). Note that for a SESOI $= 0.1$ the sample size was capped at a maximum of 200. Sample sizes based on an exploratory effect size in the Standard trajectory are small ($n_{rep} = 8.08 \pm 4.05$). Sample sizes based on an exploratory effect size are increased when the decision to proceed to confirmation was based on a SESOI (SESOI-Standard 0.1, 0.3, 0.5, 0.7: $n_{rep} = 61.58 \pm 71.46$, SESOI-Standard 1.0: $n_{rep} = 45.89 \pm 57.17$). Error bars represent standard deviations. Plotted for $n_{init} = 10$.

#### Supplementary Figure 9

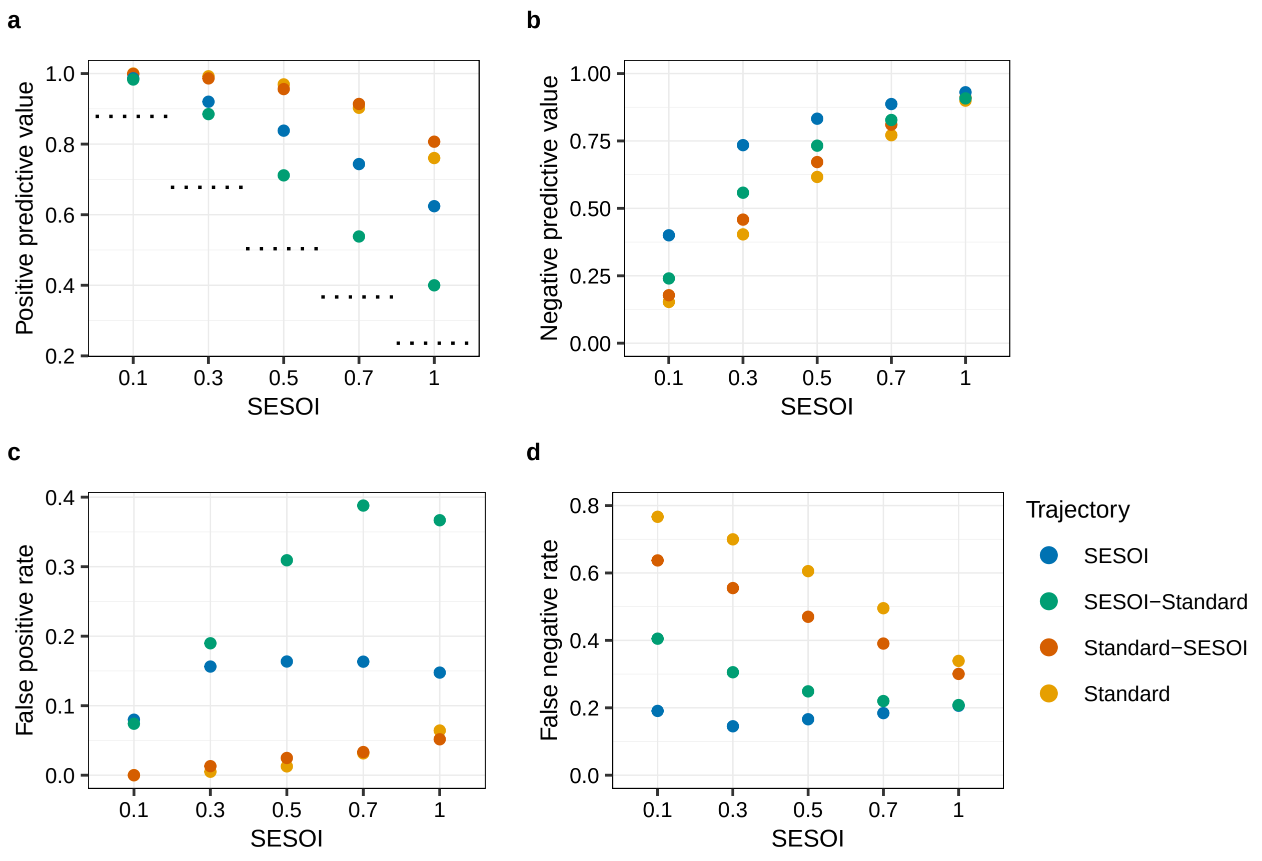


Figure S9: **Outcomes across four trajectories.** **a**, The PPV across the trajectory is consistently higher in trajectories using a significance threshold of 0.05 as decision criterion after exploration. **b**, The NPV across the trajectory is highest in SESOI trajectory. It increases with increasing SESOI across all four trajectories. **c**, The FPR is consistently higher in trajectories using a SESOI as decision criterion after exploration compared to a significance threshold of 0.05. The combination of a lenient decision criterion and low sample size in replication (SESOI-Standard trajectory) increases FPR dramatically. **d**, Trajectories using a significance threshold of 0.05 as decision criterion after exploration have a considerably higher FNR compared to trajectories using a SESOI. However, the discrepancy decreases as SESOI increases. Plotted for $n_{init} = 10$.

#### Supplementary Figure 10

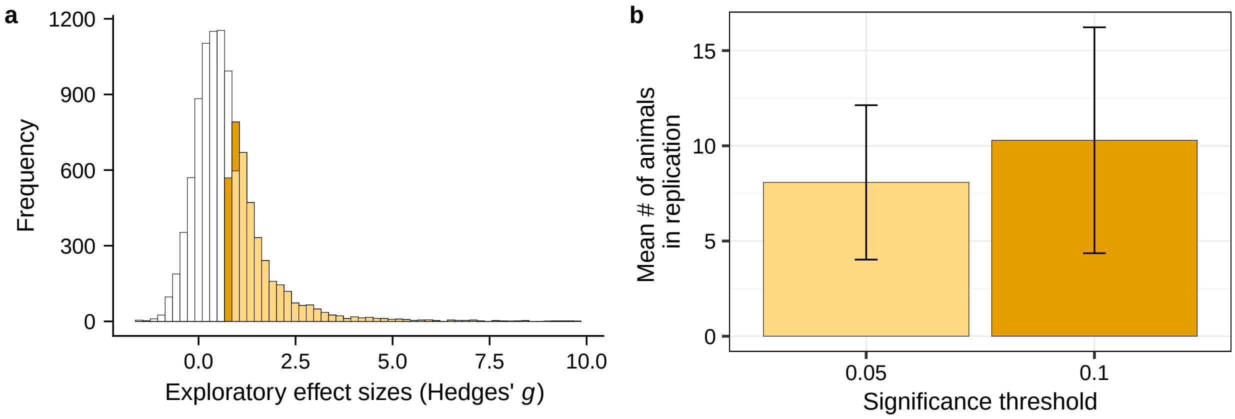


Figure S10: **Transition rates and sample sizes in confirmation for two different significance thresholds.** **a**, A significance threshold of $\alpha = 0.1$ results in a higher number of studies that transition to confirmation ($n = 4035$ compared to $n = 3272$ when $\alpha = 0.05$). **b**, A relaxed significance threshold leads to an increase in animals needed in confirmation ($\alpha = 0.1$: $n_{rep} = 10.29 \pm 5.93$, $\alpha = 0.05$: $n_{rep} = 8.08 \pm 4.05$). This is due to smaller exploratory effect sizes that are selected for confirmation when the significance threshold is less strict. Error bars represent standard deviations. Plotted for $n_{init} = 10$.

#### Supplementary Figure 11

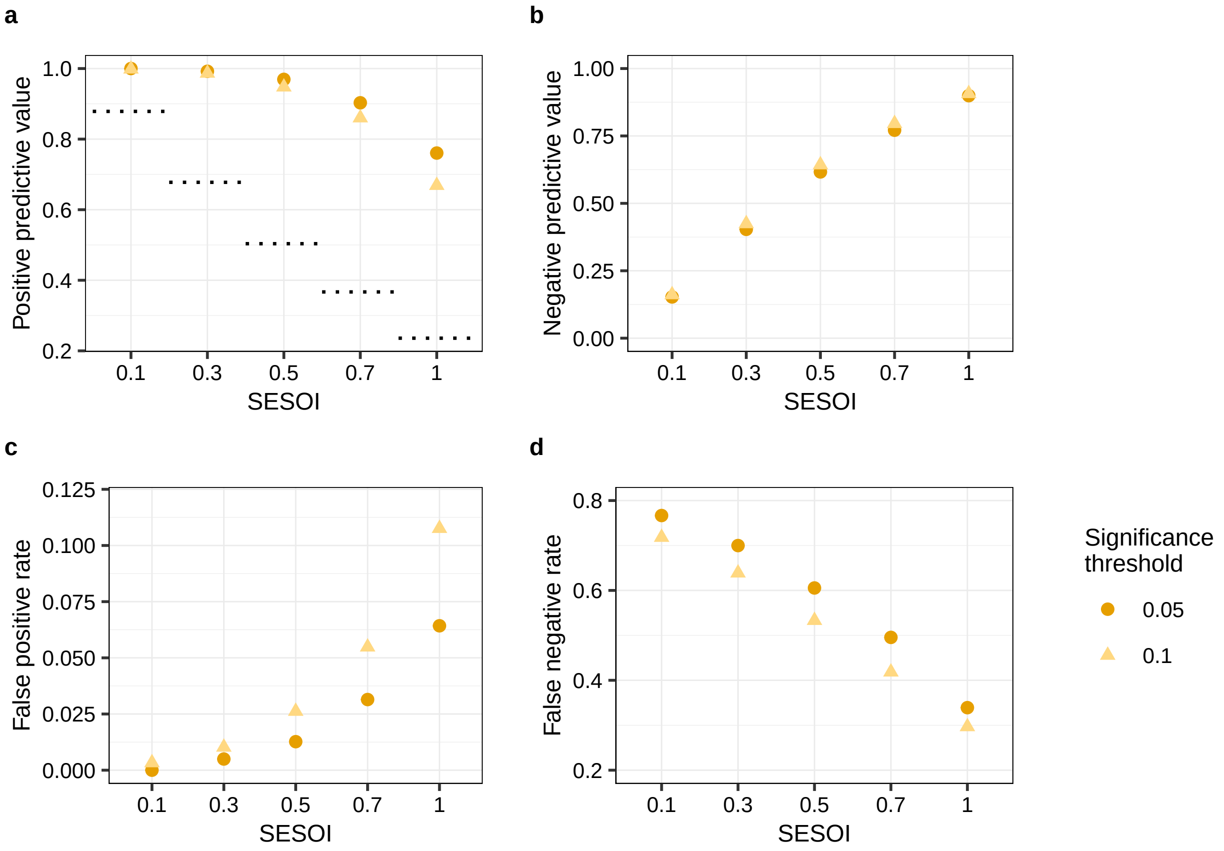


Figure S11: **Outcomes across Standard trajectory for two different significance thresholds.** **a**, The PPV across trajectory is consistently above the prior probability (dotted lines) for both significance thresholds. A relaxed significance threshold decreases the PPV for larger SESOI. **b**, A relaxation of the significance threshold does not appreciably impact the NPV. **c**, As SESOI increases the overall FPR as well as difference between FPR for $\alpha = 0.05$ and $\alpha = 0.1$ increases. **d**, The FNR is consistently higher for $\alpha = 0.05$ across SESOI, as more relevant effects are missed when a more conservative significance threshold is used. Plotted for $n_{init} = 10$.

#### Supplementary Figure 12

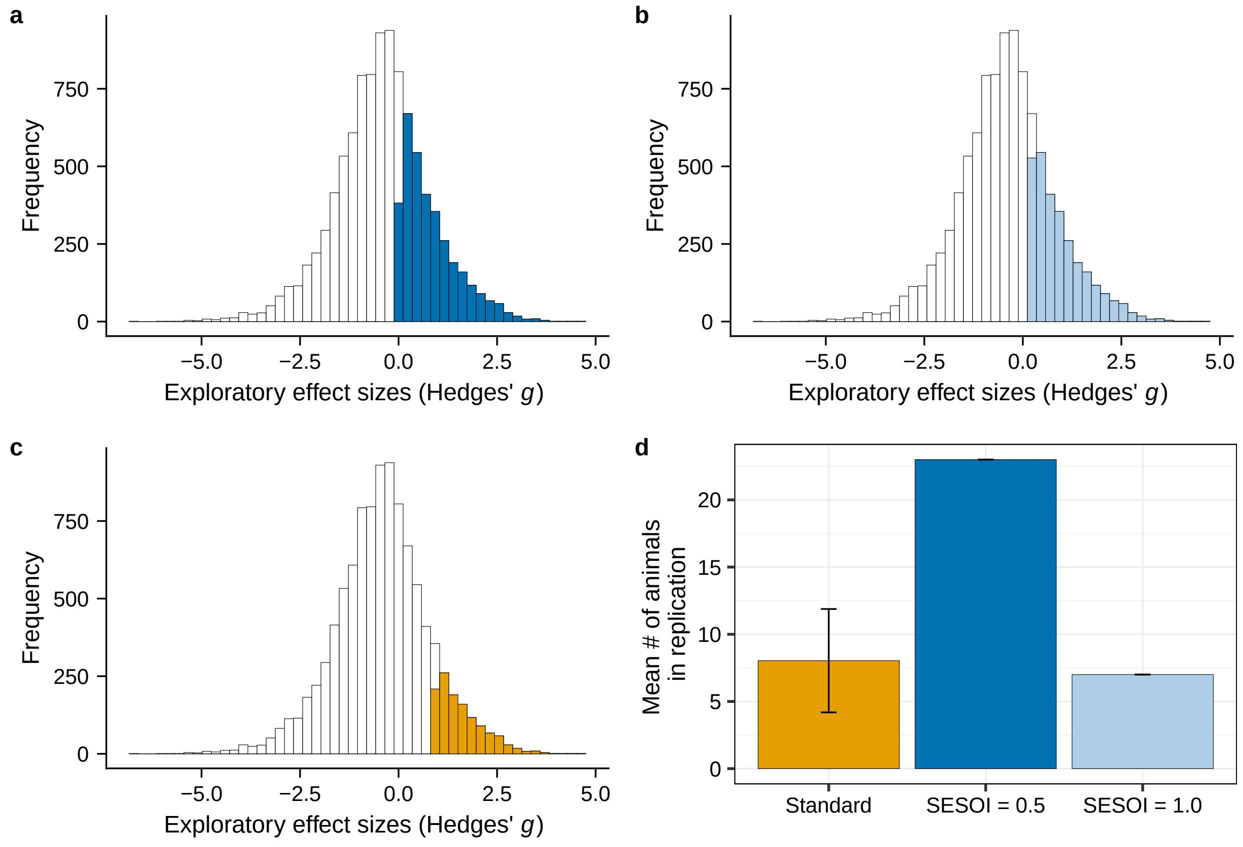


Figure S12: **Transition rates and sample sizes in confirmation for pessimistic effect size distribution. a-c**, Exploratory effect sizes ($n$ = 10000). The shaded bars show the effect sizes that were identified for confirmation using one of the two decision criteria. **a**, Dark blue shaded bars indicate effect sizes that were selected using a SESOI of 0.5 ($n$ = 3377 [1672 $>$ 0.5]). **b**, Light blue shaded bars indicate effect sizes that were selected using a SESOI of 1.0 ($n$ = 2852 [990 $>$ 1.0]). **c**, Yellow shaded bars indicate those effect sizes that were identified for confirmation using a significance threshold of 0.05 ($n$ = 1224 [1100 $>$ 0.5; 860 $>$ 1.0]). **d**, Number of animals needed in the confirmatory study. In the Standard trajectory, sample sizes are low ($n_{rep}$ = 8.03 $\pm$ 3.85), as they are based on large exploratory effect sizes. Error bars represent standard deviations. In case of trajectories using a SESOI, the number of animals is fixed. Using a SESOI of 0.5 results in $n_{rep}$ = 23, whereas a SESOI of 1.0 achieves a sample size comparable to those in the Standard trajectory ($n_{rep}$ = 7). All sample sizes displayed and reported in the text are the number of animals needed in *each* group (control and intervention).
